# ShareLoc – an open platform for sharing localization microscopy data

**DOI:** 10.1101/2021.09.08.459385

**Authors:** Jiachuan Bai, Wei Ouyang, Manish Kumar Singh, Christophe Leterrier, Paul Barthelemy, Samuel F.H. Barnett, Teresa Klein, Markus Sauer, Pakorn Kanchanawong, Nicolas Bourg, Mickael M. Cohen, Benoît Lelandais, Christophe Zimmer

## Abstract

Novel insights and more powerful analytical tools can emerge from the reanalysis of existing data sets, especially via machine learning methods. Despite the widespread use of single molecule localization microscopy (SMLM) for super-resolution bioimaging, the underlying data are often not publicly accessible. We developed ShareLoc (https://shareloc.xyz), an open platform designed to enable sharing, easy visualization and reanalysis of SMLM data. We discuss its features and show how data sharing can improve the performance and robustness of SMLM image reconstruction by deep learning.

---

Single molecule localization microscopy (SMLM) has matured to one of the most popular super-resolution methods, and has been applied to a wide range of questions across many areas of biology^1^. This success has inspired the development of a host of analysis techniques to extract localizations from raw images or infer biologically meaningful quantities from extracted localizations^1–4^. The development of further analytical techniques could greatly benefit from easy access to SMLM data generated by the community. This is especially true for recent machine learning approaches such as deep learning, whose performance is expected to increase with the amount of training data^5–8^. However, despite the large number of studies using SMLM^1^, the overwhelming majority of SMLM data remains inaccessible to the community. Although several initiatives to share imaging and other types of data already exist, such as Figshare, Zenodo, or IDR^9,10^, these are generic in purpose and are not optimally suited to gather and exploit SMLM data in a manner consistent with the FAIR principles (findable, accessible, interoperable, reusable)^11^. An important specificity of SMLM data is that each super-resolution image is built by extracting molecular coordinates from many thousands of low-resolution images (**Fig. 1a**), which often reach cumulated sizes of many GB-this is further exacerbated by a trend towards faster and longer imaging, or extensions to high-throughput or volumetric imaging^1,12–15^. Extracted localization files, though much smaller than the raw data, are usually too large to be sent by e-mail, and come in various formats, complicating reanalysis, and metadata are often unstructured or simply absent. Another restriction of generic data repositories is that they lack specific tools to visualize SMLM data. As a consequence of these limitations, only a minuscule fraction of the acquired SMLM data is made easily and publicly available, preventing their reanalysis and slowing the progress of machine learning methods dedicated to SMLM. Here, we present an open platform called ShareLoc designed to facilitate the sharing, visualization, and community-based reutilization of SMLM data.

**Figure 1:**
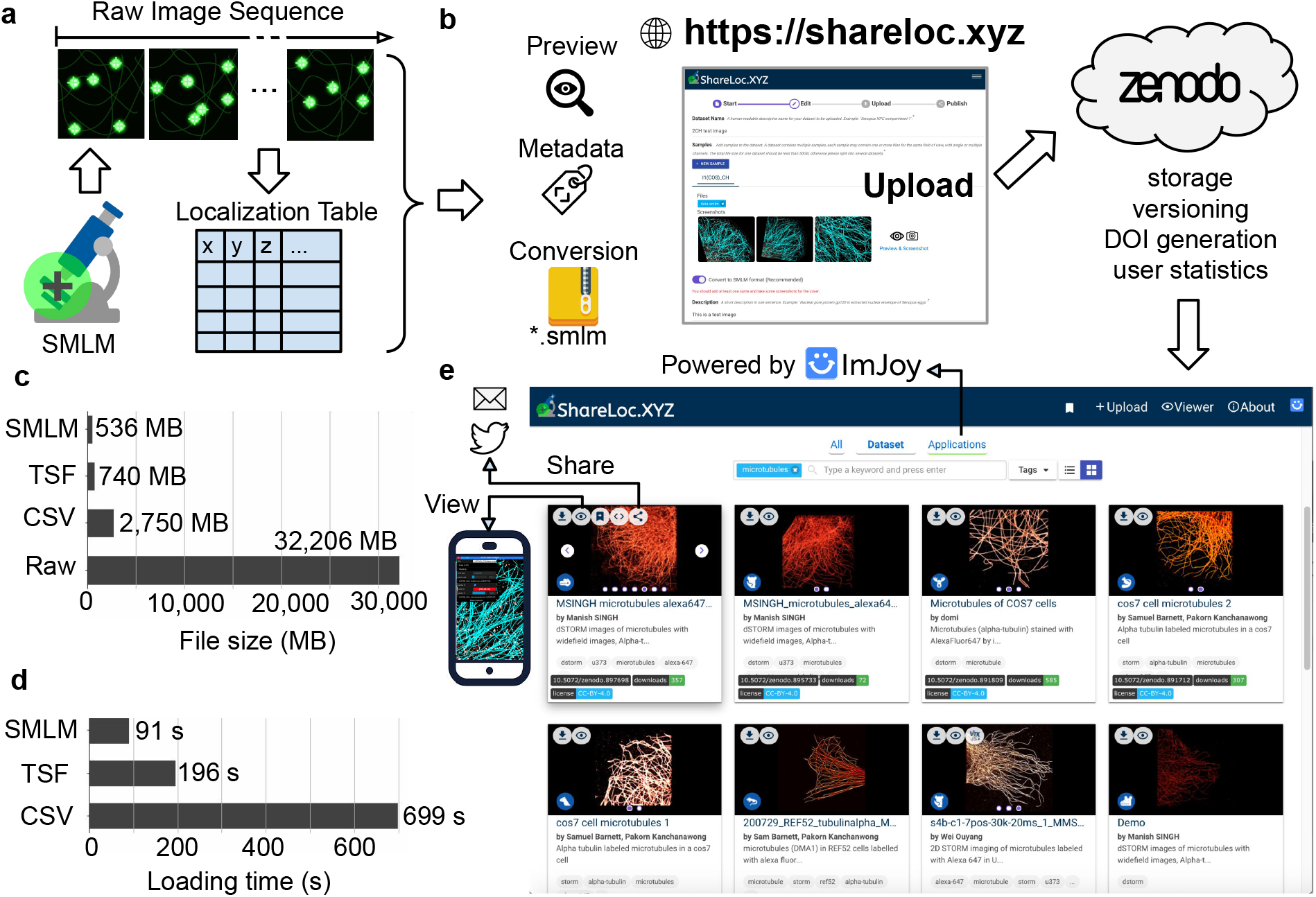
ShareLoc facilitates sharing, reanalysis and visualization of SMLM data. **a**) In SMLM, a sequence of raw, low resolution (diffraction limited) images of single, photoswitching, fluorescent molecules is acquired and their localizations are extracted using one of many localization software^1,23^, typically resulting in a localization table. **b**) The online ShareLoc platform (shareloc.xyz) allows cloud storage of SMLM data on Zenodo, and enables easy annotation of data with relevant metadata, as well as conversion to *.smlm format. **c**) The *.smlm file format provides reduced storage size for SMLM data compared to the common formats CSV and TSF and to the raw image data. **d**) The reduced file size allows corresponding gains in loading time. **e**) ShareLoc provides gallery views of uploaded data, searchable by tags, enables quick data sharing by e-mail or social media and easy visualization (2D or 3D) through a dedicated ImJoy plugin, that runs in a web browser, on computers or mobile devices. Data sets can be connected to other applications powered by ImJoy, allowing additional visualization or analysis capabilities.

The ShareLoc platform (**Fig. 1b**) consists of two parts: (i) a storage service backed by Zenodo, a widely-used general-purpose open-access repository, and (ii) an extendable system of web plugins built on top of ImJoy^16^, a platform for building and deploying interactive data science tools, that now comprises dedicated plugins for SMLM visualization. Using a Zenodo login, users can easily upload and store SMLM data sets up to 50 GB each (including localization data and/or raw images) through the online ShareLoc platform (https://shareloc.xyz), and a digital object identifier (DOI) will be automatically generated for future reference. Upon approval by the ShareLoc team, the new data will be shown on the ShareLoc repository and available for download, export, and reanalysis (**Fig. 1e**). Each data set can be linked to other data sets or analysis tools, and by default SMLM data will be linked to the dedicated SMLM visualization plugin (see below).

To facilitate rapid transmission and sharing of SMLM data, we developed a new dedicated compressed binary file format (extension *.smlm). The *.smlm data format allows significant reductions to the size of localization files, e.g. by more than three-fold compared to CSV (comma-separated values) text files and by 27% compared to TSF (tagged spot file, another binary format), without any loss of information; compared to the raw images, the size reduction is roughly fifty-fold (**Fig. 1c, Supplementary Figure 1a**). This allows comparable reductions in transfer and loading times even when accounting for file decompression delays (**Fig. 1d, Supplementary Figure 1b**). The simple binary encoding and a standard zip implementation are ideally suited for resource-constrained browser environments. The *.smlm format is portable and compact, as it can store localization coordinates and metadata (such as identifiers of the imaged molecules, antibodies, dyes, or microscopy parameters). Our file format is also flexible, as it allows easy extensions to, for example, 3D or multicolour SMLM data or tiling multiple fields of view (**Supplementary Note 1**). The ShareLoc platform allows exporting SMLM data as CSV files, if needed, and we release an ImageJ plugin to import and export *.smlm files in conjunction with ThunderSTORM, a widely used localization software^17^ (**Supplementary Software 1**), as well as command line tools in Python allowing batch downloading and conversion of single or multiple data sets^18^.

Taking advantage of this simple and efficient file format, we developed a WebGL-based viewer plugin in ImJoy^16^, for quick and fluid visualization of 2D, multicolour or 3D^19^ SMLM images. This viewer includes zooming, panning and rotation features (**Fig. 1e** and **Supplementary Movie 1**). Unlike previous SMLM visualization software^17,20,21^, the viewer plugin is entirely browser-based, requires no installations or updates and operates across all operating systems, including mobile devices.

The SMLM viewer is integrated (along with other plugins) in the online ShareLoc platform and makes it straightforward to visualize SMLM data uploaded onto Zenodo using various formats (including *.smlm, *.csv and *.xls). Data on ShareLoc can be easily shared, for example as a simple link by e-mail or using social networking platforms like Twitter (**Fig. 1e**). ShareLoc is already used by multiple labs and contains a public repository of over a hundred data sets, including images of microtubules, actin filaments, nuclear pores, clathrin coated pits and mitochondria, which can be easily browsed from a “gallery” view (**Fig. 1e, Supplementary Figure 2**). A simple and flexible tagging system is used to annotate individual images and search for particular tags, such as “microtubules” or “dna-paint” or “3d” (**Supplementary Note 2**).

Large image data sets are a particularly valuable resource for deep learning, where performance and robustness increase with the quantity and variety of training data^22^. Retraining deep models on image data sets obtained by different teams under different conditions is especially important to avoid over fitting on irrelevant image features and to enable optimal performance (e.g. image classification accuracy) on similar image data obtained by third parties. We previously developed ANNA-PALM, a deep learning method that can reconstruct high quality super-resolution images from rapidly acquired, sparse, SMLM data, after being trained on high quality super-resolution images obtained with long acquisition times^6^. Our initial demonstration of ANNA-PALM on microtubules was based on training a neural network on SMLM data from only seven fields of view imaged in our lab (totalling 372,874 individual frames and ∼161 million localizations). ShareLoc now provides access to SMLM images of microtubules from 182 fields of view, imaged by five different teams, totaling data from over 8 million individual frames and over 1 billion localizations. These images were obtained using different imaging protocols, that included differences in labelling methods (e.g. different antibodies and/or different dyes), fixation technique, cameras, objectives, laser excitation and localization software (**Supplementary Table 1, Supplementary Figure 3**). This provides the opportunity to train ANNA-PALM on a larger and more diverse data set, potentially leading to better reconstruction quality and robustness. We wished to separately address the benefit of increased data quantity vs increased diversity, and therefore first compared the quality of ANNA-PALM reconstructions after training on one vs. 60 images obtained from a single lab (lab Z), using a separate sparse localization image (and corresponding widefield image) from the same lab as test data (**Supplementary Figure 4**). Reconstruction quality improved significantly when training on 60 instead of one image, as shown by visual inspection (**Supplementary Fig. 4c,d,e)** and confirmed by multi-scale structural similarity index (MS-SSIM) analysis (p=2×10^−4^, n=21 different test images) (**Supplementary Fig. 4f**). Next, we compared ANNA-PALM reconstruction quality after training on seven images from four different labs (A,L,Z,S) vs. seven reference images from lab Z (the same images used to train our original ANNA-PALM model^6^) and testing on images from a fifth lab (lab K) (**Supplementary Fig. 5**). Again, reconstruction quality improved, both upon visual inspection and as measured by intensity profiles and MS-SSIM analysis (p<5×10^−3^, n=17 test images) (**Supplementary Fig. 5c-g**). We repeated this analysis by rotating the training and testing data sets among the five labs three more times (cross-validation). For two of these three rotations, reconstruction quality improved very significantly (p<3×10^−3^, n=19 or 44), whereas for the remaining rotation the difference was not significant (p=0.3), likely owing to the smaller number of test images (n=8) (**Supplementary Figure 5f,g**). Overall, this analysis suggests that increased training data diversity is beneficial even at constant training data size. Finally, we trained ANNA-PALM on all available 78 images from four labs (A,L,Z,S) and applied this model to reconstruct test images from a fifth lab (lab K), consisting of paired widefield and sparse localization data (**Figure 2a,b**). While the reference ANNA-PALM model trained on the seven reference images from lab Z (M0) performed well, some reconstructions exhibited imperfections, as illustrated in **Figure 2c,e**. The ANNA-PALM model trained on 78 images from four labs, however, corrected many of these imperfections (**Figure 2d,e**). We again repeated the analysis by training four ANNA-PALM models on different subsets of the data and testing on data from the remaining lab (**Figure 2f,g**). For three out of four data rotations, ANNA-PALM reconstructions had significantly higher quality compared to our reference model M0 (model M1: p<2×10^−3^, n=19; model M2: p<0.02, n=44; model M4: p=5×10^−4^, n=17) (**Figure 2g**). For model M3, there was no significant difference in reconstruction quality (p=0.89, n=8), again likely because of the small size of this test data set (n=8). Overall, these results indicate that, in agreement with our expectation, ANNA-PALM reconstructions generally improve when the model is trained on a larger and more diverse set of images. Importantly, we showed this on test images obtained from a different lab than the training data, indicating the method’s robustness to changes in microscopy parameters.

**Figure 2:**
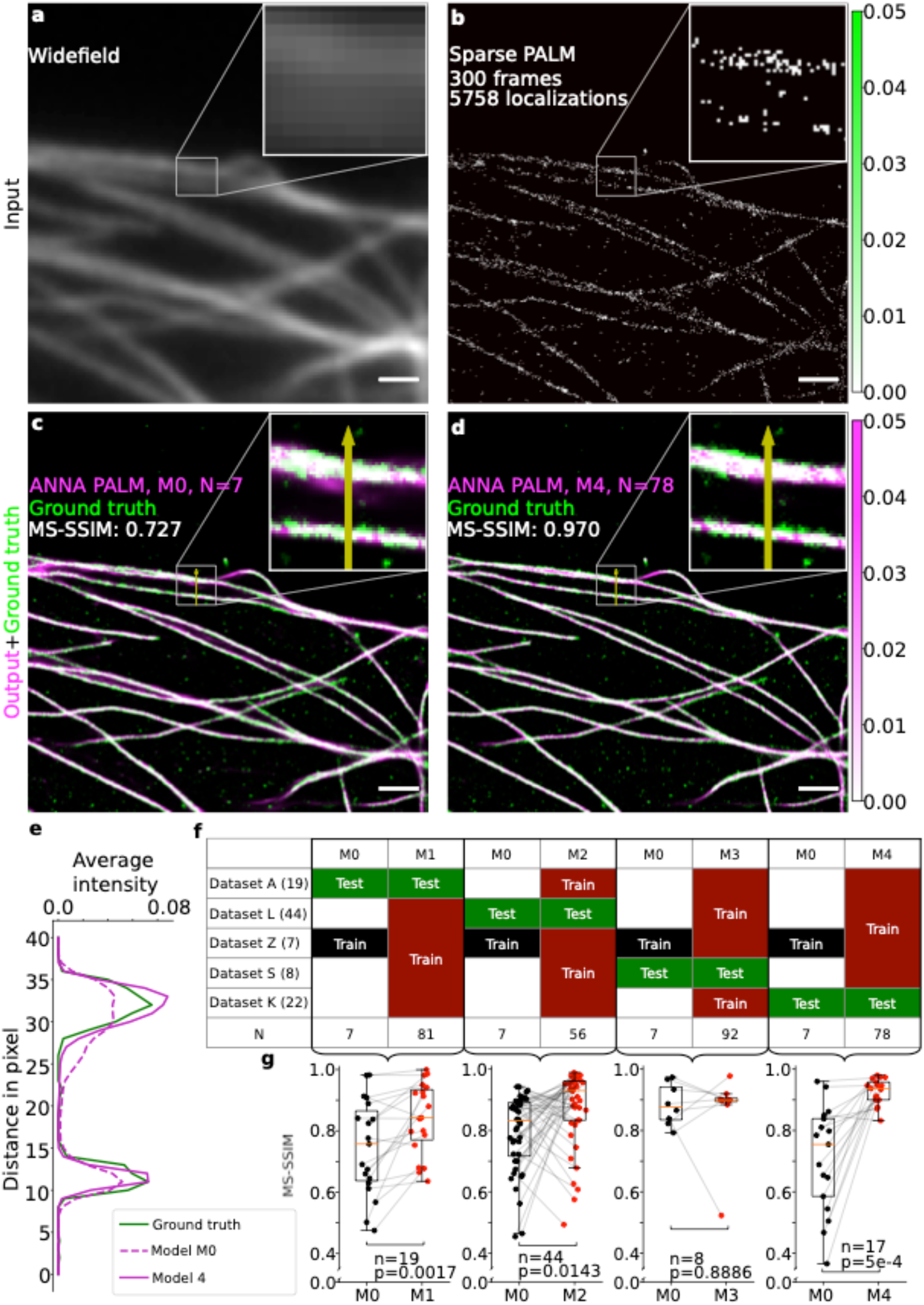
Deep learning-based image reconstruction improves when training on larger and more diverse data. **a,b**) Input images. A widefield image (**a**) and a sparse localization image (**b**) of immunolabeled microtubules. The sparse localization image has 5,758 localizations and is obtained from a sequence of 300 frames (**b**). **c,d**) Output images of two ANNA-PALM models (pink) shown in comparison to the ground truth SMLM image obtained from 20,000 frames (green). Model M0 (**c**) was trained on seven images from lab Z. Model M4 (**d**) was trained on 78 images from labs A, L, Z, and S. MS-SSIM, multi-scale structural similarity index. Scale bar in **a-d**: 1 µm. **e**) Intensity profiles of the two ANNA-PALM images and the ground-truth image along the yellow rectangle with arrowhead shown in the insets of **c,d**. The intensity peak on the bottom is well recovered by both models; for the top peak, the width predicted by model M0 is too large, but is well predicted by model M4. **f,g**) Quantitative comparisons of ANNA-PALM reconstruction quality using models trained on different data subsets. **f**) Overview of data sets used for training and testing in the four experiments. Model M0 was trained on the same seven reference images from lab Z as the previously described model^6^ and tested on images from each of the four other labs, in turn. Models M1-M4 were trained on images from four labs in different combinations, and tested on the images from the remaining fifth lab, as indicated. The number of training images, N, is indicated for each case. **g**) Boxplots compare the MS-SSIM of ANNA-PALM reconstructions relative to the ground truth using model M0 or models M1 to M4, on the same test data. Each dot corresponds to a single test image. The number of test images n is indicated. Medians are shown as horizontal red lines. Grey lines link data points corresponding to the same image. Indicated p-values are from Wilcoxon signed-rank tests. Note that the median reconstruction quality improves for all four models M1-M4 compared to M0. The increase is significant for all models except M3.

The ShareLoc platform addresses all four principles of FAIRness proposed for scientific data, and which are presently lacking for SMLM. Findability is enabled by ShareLoc’s data annotation and searching features. Accessibility is facilitated by the compact *.smlm file format and the simple and fluid viewer. Interoperability is ensured by the open source format and tools to import and export SMLM data into and out of the ShareLoc platform. Reusability is made easy by the use of a unified file format and structured metadata. We illustrated this by showing how reanalysis of data gathered on ShareLoc can improve the quality and robustness of an advanced image reconstruction method based on machine learning. More generally, we believe that ShareLoc will accelerate the sharing and development of new analytical methods for SMLM, the mining of existing SMLM data for new biological information, and will help promote reproducible research in single molecule imaging and its numerous applications in life science research.

## Supporting information

Supplementary Software 1

## Acknowledgements

We thank Felix Woitzel for development and sharing of the Fairy Dust viewer. We thank M. Lelek for advice on metadata. CL acknowledges funding from CNRS ATIP AO2016. SB and PK acknowledge funding support from Mechanobiology Institute seed funding, Singapore Ministry of Education (MOE2019-T2-2-014) and National Research Foundation (QEP-P7). MS, MC and CZ acknowledge funding by Agence Nationale de la Recherche (grant ANR 17 CE13 0026 02), and MC by Labex DYNAMO (ANR-11-LABX-0011-DYNAMO) and MITOFUSION (ANR-19-CE11-0018). JB, BL and CZ acknowledge funding by Institut Pasteur and the Region Ile-de-France in the framework of DIM ELICIT.

