## Supplement for "ShareLoc – an open platform for sharing localization microscopy data"

### Shareloc – a web platform to share and visualize localization microscopy data

**Supplementary Note 1: The *.smlm data format**

This Note details our motivation for developing the *.smlm data format and its design principles.

Storing and transmitting the data underlying typical SMLM experiments is challenging, because of the enormous file size. A typical experiment requires a sequence of ~$10^4$-$10^5$ individual raw (diffraction limited) images for each super-resolution image and data volume can reach ~100 GB per experiment or more[1]. With fast, kHz-rate cameras, a SMLM system can output Terabytes per hour. The localization data computed from the raw images are much smaller, but still typically occupy several GB, because by default, localizations are generally stored as comma separated value (CSV) files, which trade off file size to optimize human readability. As a result, typical SMLM data files are too large for simple sharing methods such as e-mail.

To address this, we designed a specific data format called 'SMLM' (*.smlm), which instead optimizes for size rather than human readability, and can reduce file size by 6-fold or more compared to the CSV format. For example, a 7.16 GB *.csv localization table can be replaced by a 1.34 GB *.smlm file without any loss of information. To achieve this size reduction, we adopted two strategies. First, we use binary encoding rather than text encoding, implying that all numbers are represented by data types such as integer, float or double. Second, we compress the binary file using generic lossless compression algorithms that come with the standard zip file format[2]. In order to compensate the lack of human readability of binary formats, we add a complementary text file (hereafter called 'manifest file') to describe the structure of the localization table and store metainformation. See an example layout of a *.smlm file below.

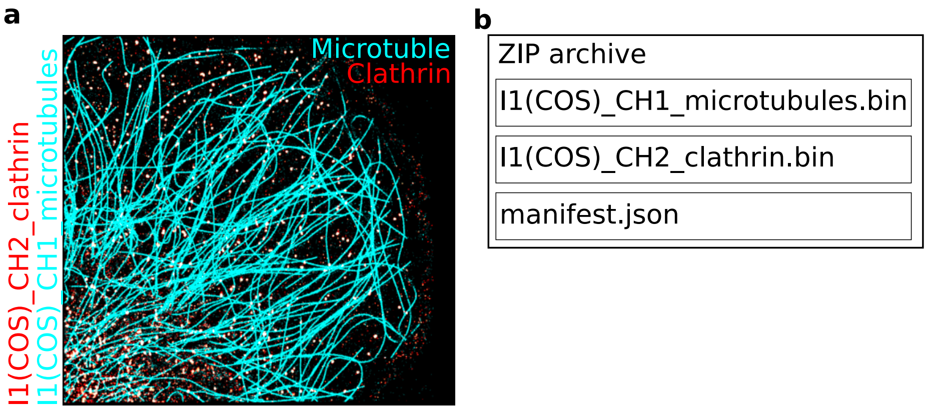

An example SMLM image (**a**) and the structure of its corresponding *.smlm file (**b**). **a**) Dual color 3D SMLM image of clathrin (red) and microtubules (cyan), from the Leterrier lab. **b**) Layout of the corresponding *.smlm file. The *.smlm file is a zip archive that contains two binary localization files (one for each color channel), and a manifest file with the fixed name ‘Manifest.json’. Note that the *.smlm format can also be used to store raw images, for example widefield images corresponding to the SMLM image.

The manifest file is implemented in the widely supported JavaScript Object Notation (*.json) format and serves two purposes. First, it stores information that is needed for a reader program to properly read the binary data, such as the data type (e.g. float or double), the number of rows, or the header. Second, it provides a human understandable summary of the localization table that does not require loading or unzipping the whole file.

```json
{
    "format_version": "0.2",
    "formats": {
        "smlm-table(binary)-0": {
            "name": "smlm-table(binary)-0",
            "type": "table",
            "mode": "binary",
            "extension": ".bin",
            "columns": 6,
            "headers": ["frame","x","y","z","uncertainty_xy","uncertainty_z_nm_"],
            "dtype": ["float32","float32","float32","float32","float32","float32"],
            "shape": [1,1,1,1,1,1],
            "units": [],
            "description": "the default smlm format for localization tables"
        },
        "smlm-table(binary)-1": {
            "name": "smlm-table(binary)-1",
            "type": "table",
            "mode": "binary",
            "extension": ".bin",
            "columns": 6,
            "headers": ["frame","x","y","z","uncertainty_xy","uncertainty_z_nm_"],
            "dtype": ["float32","float32","float32","float32","float32","float32"],
            "shape": [1,1,1,1,1,1],
            "units": [],
            "description": "the default smlm format for localization tables"
        }
    },
    "files": [
        {
            "name": "I1(COS)_CH2_clathrin.bin",
            "format": "smlm-table(binary)-0",
            "channel": "default",
            "rows": 382733,
            "offset": {
                "x": 0,
                "y": 0
            },
            "metadata": {},
            "exposure": -1,
            "type": "table",
            "min": {
                "frame": 1,
                "x": 1560.800048828125,
                "y": 1531.4000244140625,
                "z": -404.3999938964844,
                "uncertainty_xy": 0.30000001192092896,
                "uncertainty_z_nm_": 0.699999988079071
            },
            "max": {
                "frame": 39999,
                "x": 40180.1015625,
                "y": 39407.1015625,
                "z": 403.5,
                "uncertainty_xy": 14.5,
                "uncertainty_z_nm_": 29
            },
            "avg": {
                "frame": 19057.150465206876,
                "x": 15180.081666312117,
                "y": 18308.978733591597,
                "z": -164.58818115778257,
                "uncertainty_xy": 3.9080617543854737,
                "uncertainty_z_nm_": 7.816527450217723
            },
            "hash": "e360f81d1f950555ba3aeff2981392dd",
            "tags": []
        },
        {
            "name": "I1(COS)_CH1_microtuble.bin",
            "format": "smlm-table(binary)-1",
            "channel": "default",
            "rows": 769747,
            "offset": {⋯
            },
            "metadata": {},
            "exposure": -1,
            "type": "table",
            "min": {⋯
            },
            "max": {⋯
            },
            "avg": {⋯
            },
            "hash": "6f3a97652596e81dcc63ae6a333630bf",
            "tags": []
        }
    ],
    "sample": "",
    "labeling": "",
    "license": "CC BY 4.0",
    "citeAs": "",
    "hash": "90c9563de5d6272ee535bd6b4220872a"
}
```

Example manifest file for the dual color SMLM image above (only excerpts are shown). The file contains several fields, including 'formats' and 'files'. The 'formats' field stores a set of formats describing the header (here: “frame”, “x”, “y”, etc.), the data type (in this case, float32 values), the shape, etc. of each column. The 'files' field specifies which format is used to read the corresponding localization file, and also includes meta information for each binary file, such as the number of rows (here: 382,733 and 769,747, corresponding to the number of localizations for clathrin and microtubules, respectively), the number of raw image frames (39,999 for clathrin), the minimum, maximum and average values of computed x,y,z coordinates, etc.

An important feature of the *.smlm format is its flexibility. Because the structure of the binary files is specified in the manifest file, it can be easily adapted to different types of applications. For example, in addition to the required columns specifying *x* and *y* coordinates, one can easily add a third column to specify the axial (*z*) coordinate in a 3D SMLM image, as in the example above[3]. This only requires changing the “headers” in the ‘formats’ field. It is similarly straightforward to add more columns, e.g. to specify additional parameters for each localization, such as the intensity, localization uncertainty, etc. It is even possible to add additional columns to store the raw pixel values of image patches centered on each localization. Another illustration of this flexibility are multicolor images. In multicolor SMLM, localizations from each color are typically stored in separate localization files. These can be stored in the same *.smlm file, by specifying their names in the ‘files’ field of the manifest file and indicating the same format in the ‘format’ field, as also shown in the above example. Similarly, a *.smlm file can store both a localization table and the corresponding widefield image (in this case, the ‘formats’ field specifies both the table format and the image format). Another related example is tiled imaging, where images from many adjacent (or partly overlapping) fields of view are assembled into a much larger effective field of view. The many localization tables corresponding to each individual field of view can be saved together in a single *.smlm file, by specifying the names of the individual localization files in the ‘files’ field. In this case, each localization file can be given an ‘offset’ field to store the spatial offset (e.g. the coordinates of the upper left corner) of each field of view.

**Supplementary Note 2: ShareLoc data tagging system**

For optimal reuse, shared SMLM data should be annotated with metadata containing all information required to reproduce the data, including the experimental conditions, image acquisition and analysis protocols[4]. To facilitate such annotations, ShareLoc provides a simple and flexible tagging system that allows to quickly reuse existing tags and/or add new tags. Example tags are provided in the Table below. The Table also shows how tags are grouped in a number of categories, such as imaging modality, cell line, fluorophore, etc. In addition to allowing quicker tagging, using predefined tags tends to limit the proliferation of different naming conventions (e.g. "Alexa 647", "Alexa fluorophore 647", "AF647" or "alexa-647" for the same fluorophore Alexa-647). Note that for the same purpose we recommend to use only lowercase names. On the ShareLoc home page, next to the search bar, tags can also be selected to filter the gallery view and search for specific data sets. For example, selecting the tags "microtubule", "alexa-647" and "nih3t3", will display only the SMLM data of microtubules imaged with Alexa-647 in NIH3T3 cells. If the existing tags are not sufficient, users can easily add custom tags to the system. These can be added to one of the predefined categories below or in the "other" category.

| **Category** | **Tag** |
|---|---|
| Modality | palm, storm, dstorm, dna-paint, paint |
| Cell line | u2os, u373, cos7, ref52, msck, hela, nih3t3, yeast, e-coli |
| Fluorophore | alexa-647, alexa-555, alexa-532, cy5, cy3b, cf680, cf660, cf568, hmsir, atto-488, mos2, mapple, dendra2, pa-gfp, pa-mcherry, dronpa, phalloidin-647, nile-red, jf-646, jf-568, pa-jf568, pa-jf646 |
| Labeling Strategy | transient-transfection, stable-knock-in, endogenous-labeling-crispr, direct-immuno-labeling, indirect-immuno-labeling, halo-tag, snap-tag, tetracysteine-tag |
| Structure | actin, microtubules, mitochondria, nuclear-pore, clathrin, vimentin, plasma membrane, intermediate filament, dna |
| Target molecule | nup133, nup96, tom22, alpha-tubulin, beta- tubulin, actin, vimentin, wga, escrt, ftsz, h2, h3, h1, polye |
| Dimension | 2d, 3d |
| Camera | em-ccd, scmos |
| Reconstruction software | thunderstorm, smap, smoiphot, zola-3d, picasso, quickpalm |
| Buffer | gluox, catalase, mea, abbelight-safe-reagent, idylle-everspark |
| Fixation | 4% pfa, 1% pfa, pfa+gluta, glyoxal, methanol |
| Other | All other customized tags |

The Table below lists the current categories and tags. Please note that this list will be automatically extended if users add new tags.

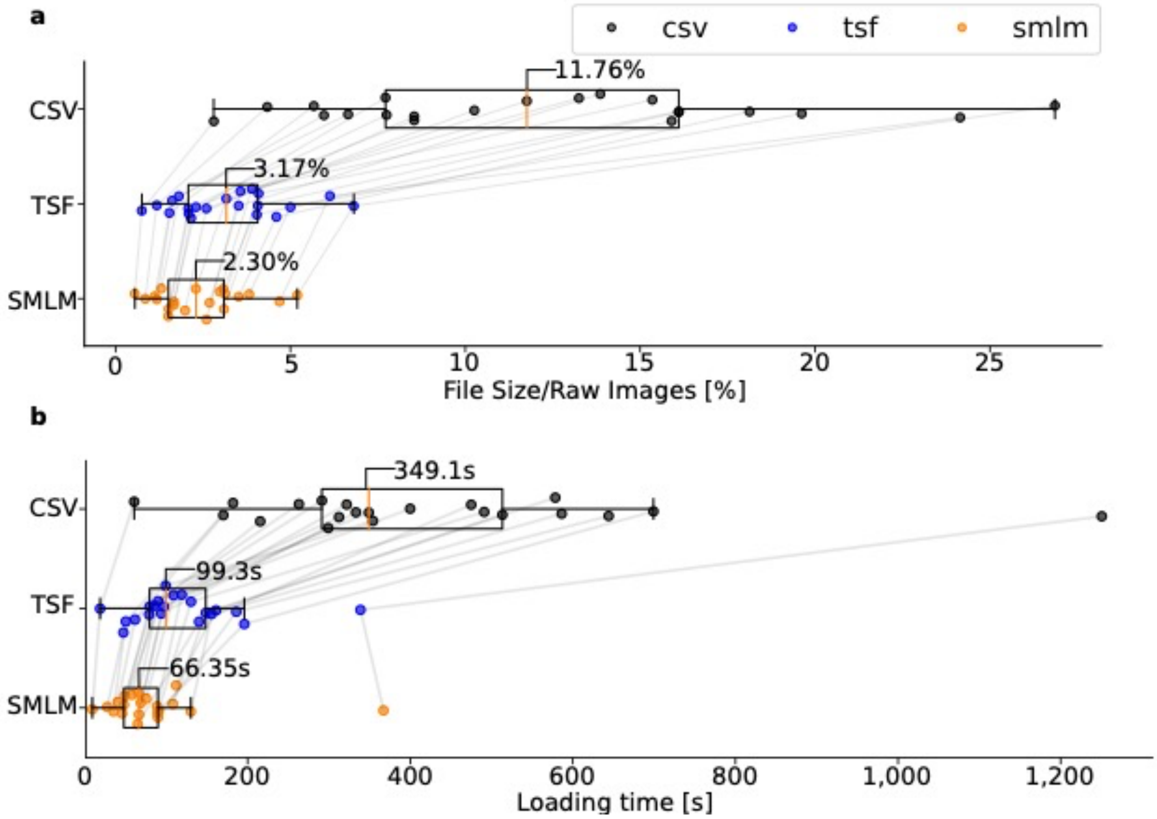

**Supplementary Figure 1: Localization data size and loading times for different file formats**

**a,b**) File sizes (**a**) and loading times (**b**) of n=21 different localization data sets, for three formats: CSV (*.csv), TSF (*.tsf) and SMLM (*.smlm). The file sizes are expressed as percentage of the raw image data size. Each dot corresponds to a distinct data set. Dots for the same data sets in different formats are connected by grey lines. Boxplots show the median as orange bars (median value is indicated), the first and third quartiles as box edges, and the full data range (except outliers) by whiskers.

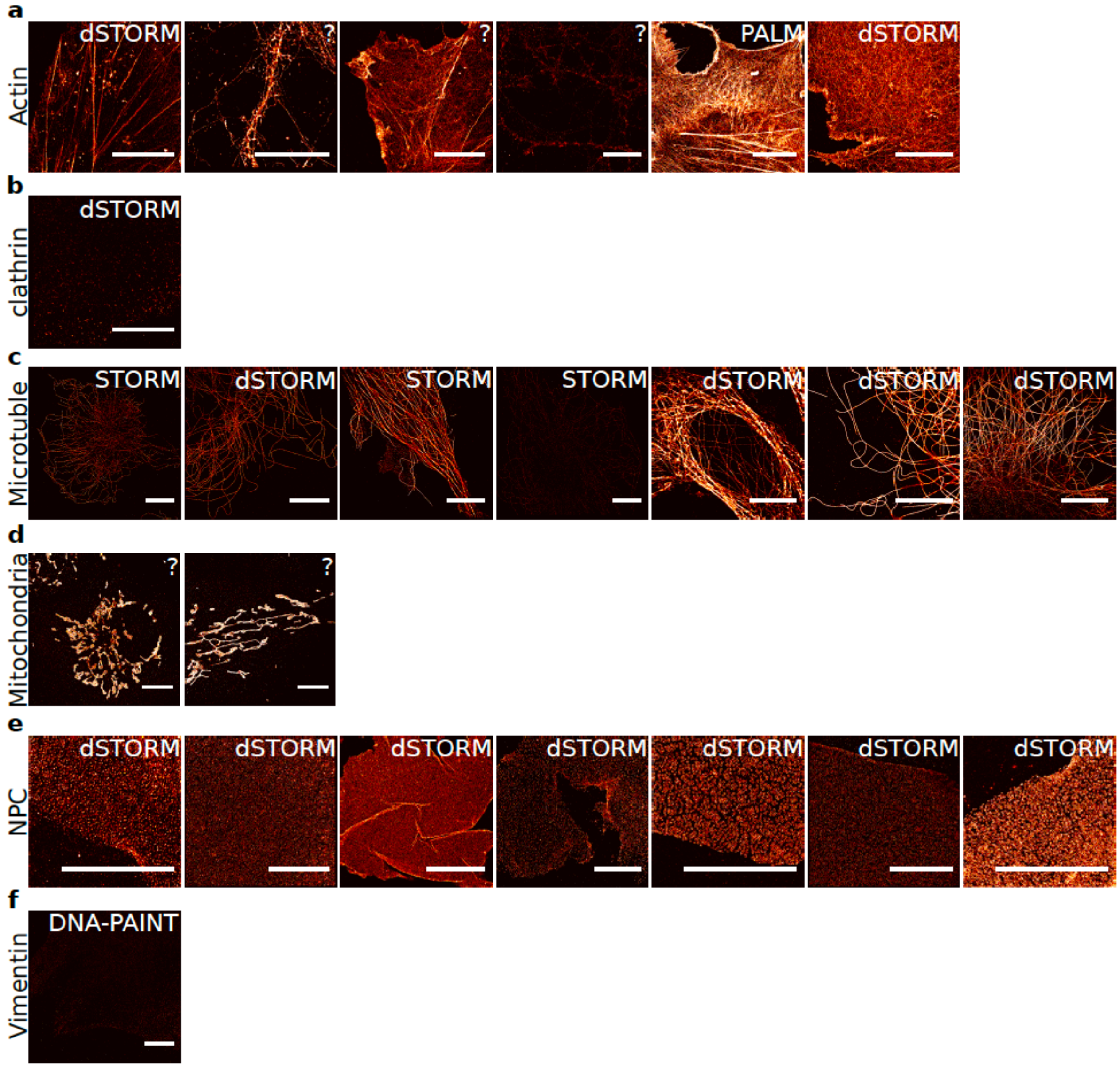

**Supplementary Figure 2: Shareloc contains diverse SMLM data sets**

This Figure shows examples of diverse SMLM data available on the Shareloc website. The data include images of actin filaments (**a**), clathrin coated pits (**b**), microtubules (**c**), mitochondria (**d**), nuclear pore complexes (NPC) (**e**), and vimentin (**f**).

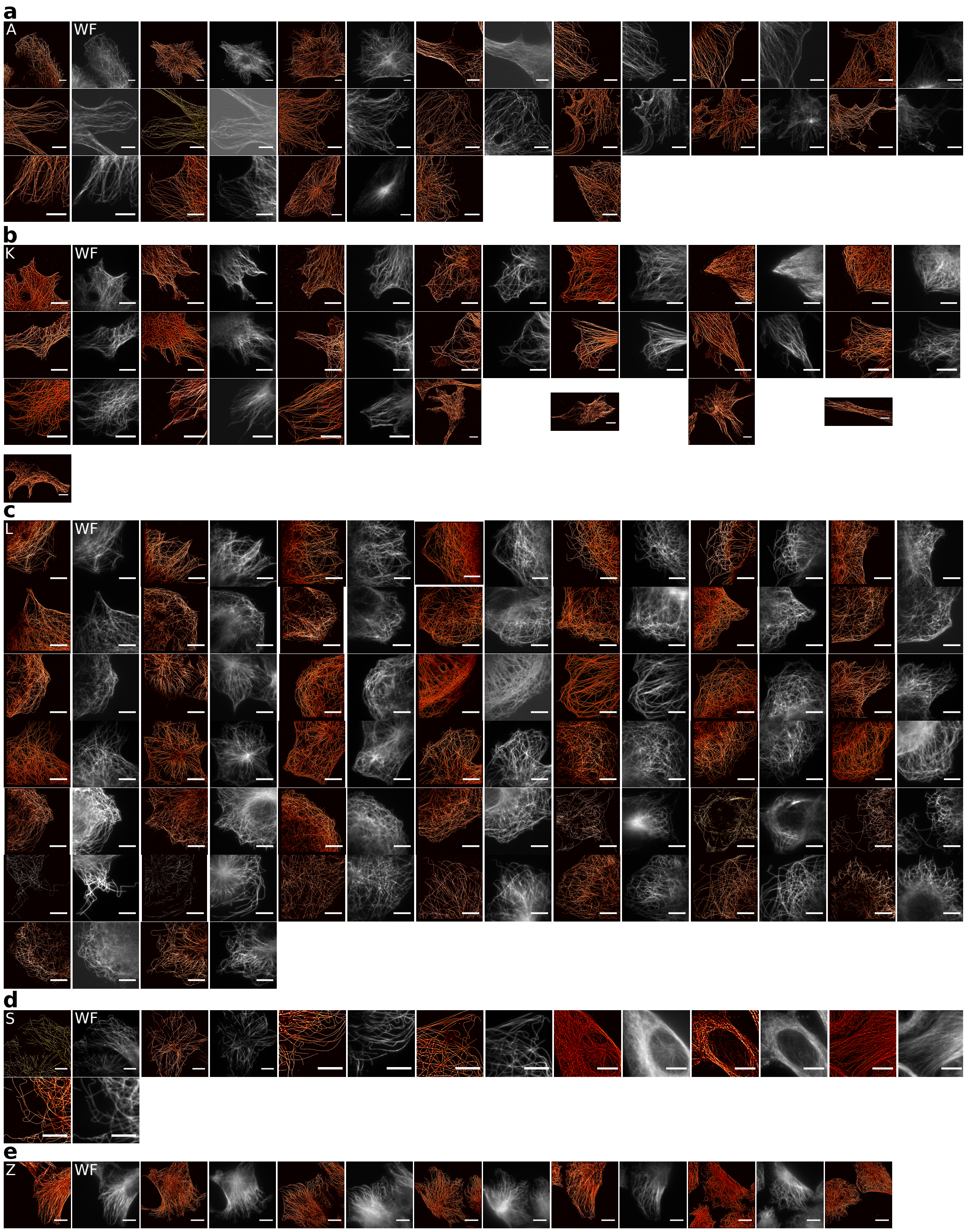

**Supplementary Figure 3: SMLM images of microtubules from multiple labs**

This Figure shows SMLM images (and for most, the corresponding widefield image) of microtubules from different labs. **a**) 19 SMLM images from lab A. **b**) 22 SMLM images from lab K. **c**) 44 SMLM images from lab L. **d**) 8 SMLM images from lab S. **e**) 7 SMLM images from lab Z. Scale bars are 10 µm.

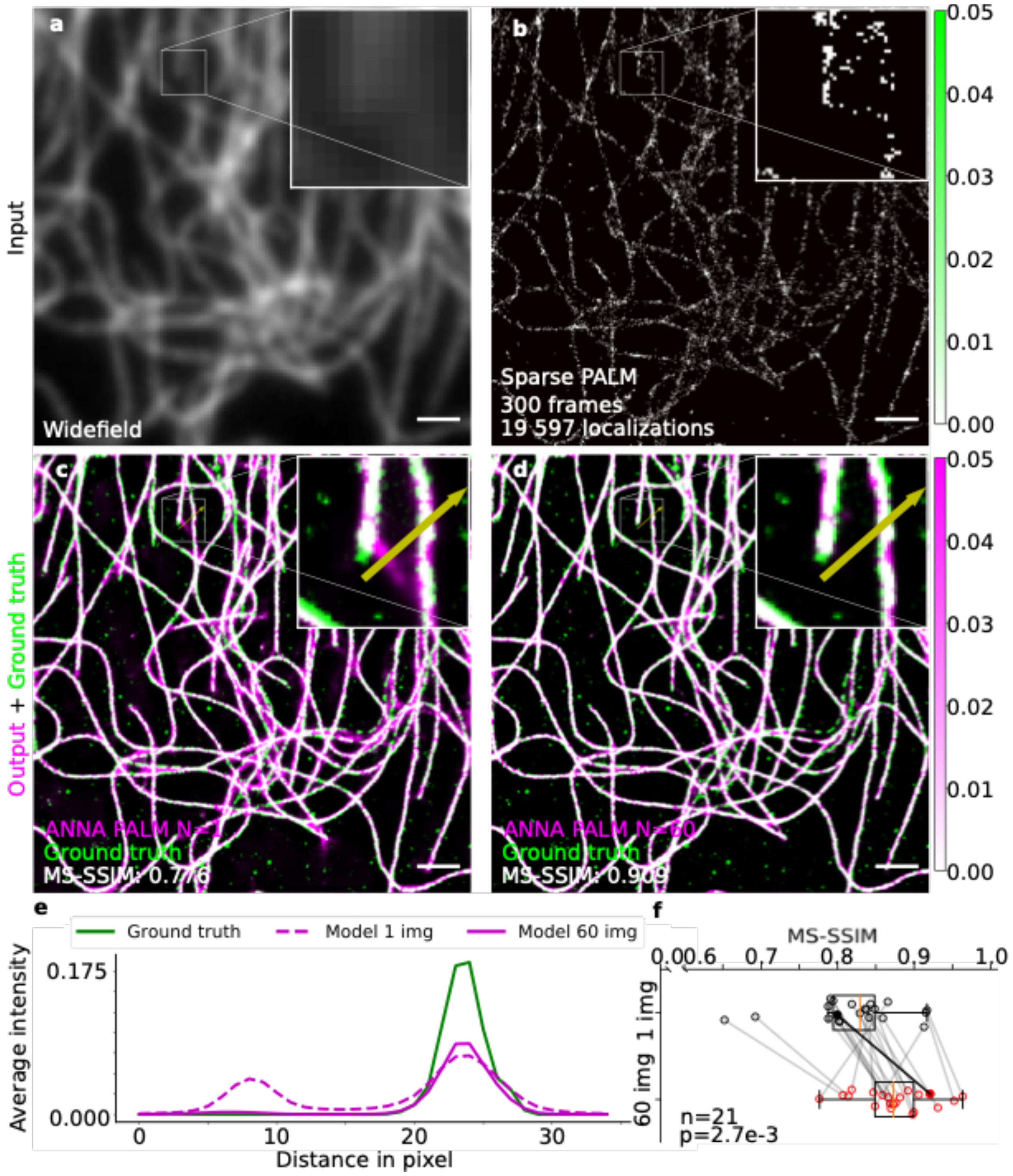

#### Supplementary Figure 4: ANNAPALM reconstruction quality increases with number of training images

**a,b**) Input images. A widefield image (**a**) and a sparse localization image (**b**) of immunolabeled microtubules. The sparse localization image is obtained from a sequence of 300 frames from lab Z (**b**). **c,d**) Output images of two ANNAPALM models (pink) shown in comparison to the ground truth SMLM image obtained from 60,000 frames (green). Model 1 (**c**) was trained on a single image (N=1) from lab Z. Model 60 (**d**) was trained on N=60 images from lab Z. MS-SSIM, multi-scale structural similarity index. **e**) Normalized intensity profiles of the two ANNAPALM images and the ground-truth image along the yellow rectangle with an arrowhead shown in the insets of **c,d**. Note that model 1 incorrectly predicts two peaks, whereas model 60 correctly recovers a single peak. **f**) Boxplot compares ANNAPALM reconstruction quality (MS-SSIM) for n=21 test images from lab Z, after training ANNAPALM on N=1 (black) vs. N=60 images (red) from lab Z. Training and testing data are entirely disjoint and together contain 81 SMLM images in total. Grey lines connect dots for identical test images. For each red dot, we trained a model on 60 other randomly chosen images from lab Z. For each black dot, we trained a model on one image, randomly chosen among the remaining 80 images. Medians are shown as vertical red lines. Indicated p-value is from a signed-rank Wilcoxon test.

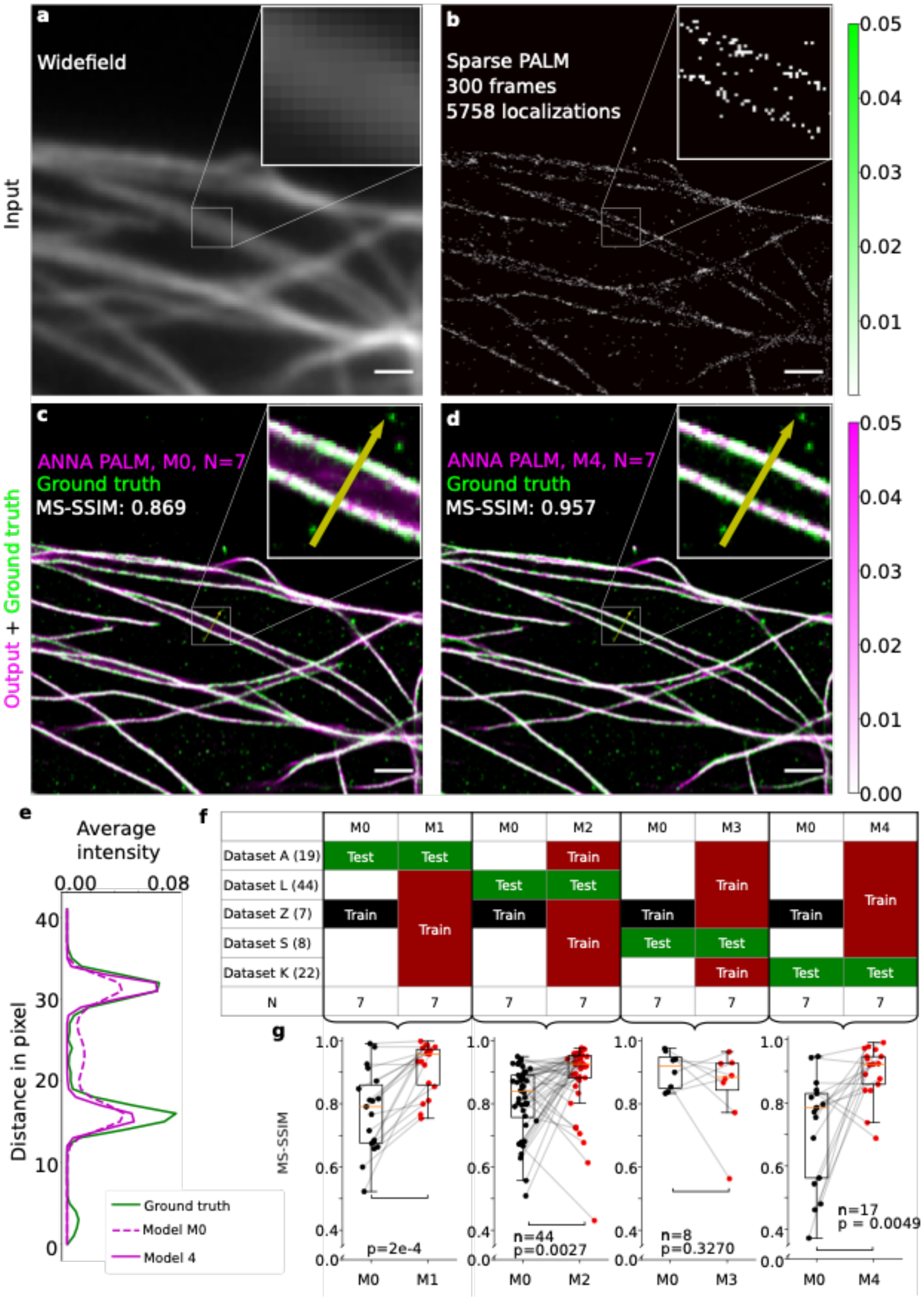

**Supplementary Figure 5: ANNAPALM reconstruction robustness increases with training image diversity**

**a,b**) Input images. A widefield image (**a**) and a sparse localization image (**b**) of immunolabeled microtubules from lab K. The sparse localization image is obtained from a sequence of 300 frames (**b**).

**c,d**) Output images of two ANNAPALM models (pink) shown in comparison to the ground truth SMLM image obtained from 20,000 frames (green). Model M0 (**c**) was trained on the same seven images from lab Z as previously[5]. Model M4 (**d**) was trained on seven images in total, originating from labs L, Z, S and K. MS-SSIM, multi-scale structural similarity index. **e**) Normalized intensity profiles of the two ANNAPALM images and the ground-truth image along the yellow rectangle with arrowhead shown in the insets of **c,d**. The two intensity peaks are well recovered by both models; but the ANNAPALM image predicted by model M0 exhibits an incorrect dimmer intermediate peak in between. This artefact is not apparent in the ANNAPALM image predicted by model M4. **f,g**) Quantitative comparisons of ANNAPALM reconstruction quality in four experiments. **f**) Overview of data sets used for training and testing in the four experiments. Model M0 (trained on images from lab Z) was tested on images from each of the four other labs, in turn. Models M1-M4 were trained on seven images chosen randomly from four labs in different combinations (with one or two images from each lab), and tested on the images from the remaining fifth lab. **g**) Boxplots compare the MS-SSIM of ANNAPALM reconstructions relative to the ground truth using model M0 or models M1-M4. Each dot corresponds to a single test image. The number of test images *n* is indicated. Medians are shown as horizontal orange lines. Grey lines correspond to the same image. Indicated p-values are from Wilcoxon signed-rank tests. Reconstruction quality improves very significantly for models M1, M2 and M4 compared to model M0.
